# Integrative spatiotemporal modeling of biomolecular processes: application to the assembly of the Nuclear Pore Complex

**DOI:** 10.1101/2024.08.06.606842

**Authors:** Andrew P. Latham, Jeremy O. B. Tempkin, Shotaro Otsuka, Wanlu Zhang, Jan Ellenberg, Andrej Sali

## Abstract

Dynamic processes involving biomolecules are essential for the function of the cell. Here, we introduce an integrative method for computing models of these processes based on multiple heterogeneous sources of information, including time-resolved experimental data and physical models of dynamic processes. We first compute integrative structure models at fixed time points and then optimally select and connect these snapshots into a series of trajectories that optimize the likelihood of both the snapshots and transitions between them. The method is demonstrated by application to the assembly process of the human Nuclear Pore Complex in the context of the reforming nuclear envelope during mitotic cell division, based on live-cell correlated electron tomography, bulk fluorescence correlation spectroscopy-calibrated quantitative live imaging, and a structural model of the fully-assembled Nuclear Pore Complex. Modeling of the assembly process improves the model precision over static integrative structure modeling alone. The method is applicable to a wide range of time-dependent systems in cell biology, and is available to the broader scientific community through an implementation in the open source *Integrative Modeling Platform* software.

## Introduction

Describing the dynamic processes involving biomacromolecules is essential for understanding the function of the cell. Thus, individual techniques, such as Förster resonance energy transfer (FRET),^1,2^ nuclear magnetic resonance,^3,4^ and physics-based molecular simulation,^5,6^ have been pioneered to characterize biomolecular processes. These methods have been successful in investigating complex phenomena, such as protein folding.^7–10^ However, describing the rearrangements of multi-component protein systems, including the process of complex assembly, remains challenging because of the scale in space and time as well as the relative lack of information.

One approach to overcoming this challenge is integrative modeling. Integrative modeling benefits from simultaneously using varied information, including from different experimental data, statistical preferences, physical theories, and other prior models.^11–13^ By maximizing the input information, integrative modeling maximizes the accuracy, precision, and completeness of the output models. Static structures of protein complexes are already routinely determined by integrative modeling.^14–22^ For example, the Nuclear Pore Complex (NPC) structure was computed by considering previously determined structures of its components, chemical cross-links between them, a cryo-electron microscopy map of the entire complex, and other information.^23–32^ The human NPC consists of *∼*1000 copies of *∼*30 different proteins, called nucleoporins (Nups). It spans the nuclear envelope and facilitates controlled macromolecular transport between the nucleus and the cytoplasm.^33–35^ However, despite the success of integrative modeling of static structures, no integrative method has yet been developed for modeling macromolecular processes, including the process of complex assembly.^36^

Here, we present and illustrate an integrative spatiotemporal modeling method. The method starts by sampling “snapshot models” at each discrete time point; a snapshot model is a set of static structural models with the same composition, where each subcomplex is assigned to a specific location in the fully assembled complex. A trajectory is composed by selecting and connecting static snapshots between neighboring time points; these trajectories are scored by their fit to the input information. This depiction of a dynamic process facilitates informing its model by the data created at discrete time points during the process. Such data are available for the assembly of the human NPC during mitosis. ^37–44^

Next, we outline the integrative spatiotemporal modeling workflow, followed by a description of our model of the NPC assembly pathway, a description of its agreement with the input information, and a comparison between our new method and an alternative approach of modeling each snapshot independently, without considering connections between the snapshots.

## Integrative spatiotemporal modeling workflow

Integrative spatiotemporal modeling of biomolecular processes is described by way of applying it to the assembly of the human NPC in the context of the nuclear envelope during cell division (SI Appendix, Table S1). We modeled the assembly process by first modeling multiple snapshots at each discrete time point (Fig. 1A), followed by connecting these snapshots into trajectory models (Fig. 1B). The output of modeling is a set of weighted trajectory models sufficiently consistent with input information.

**Figure 1:**
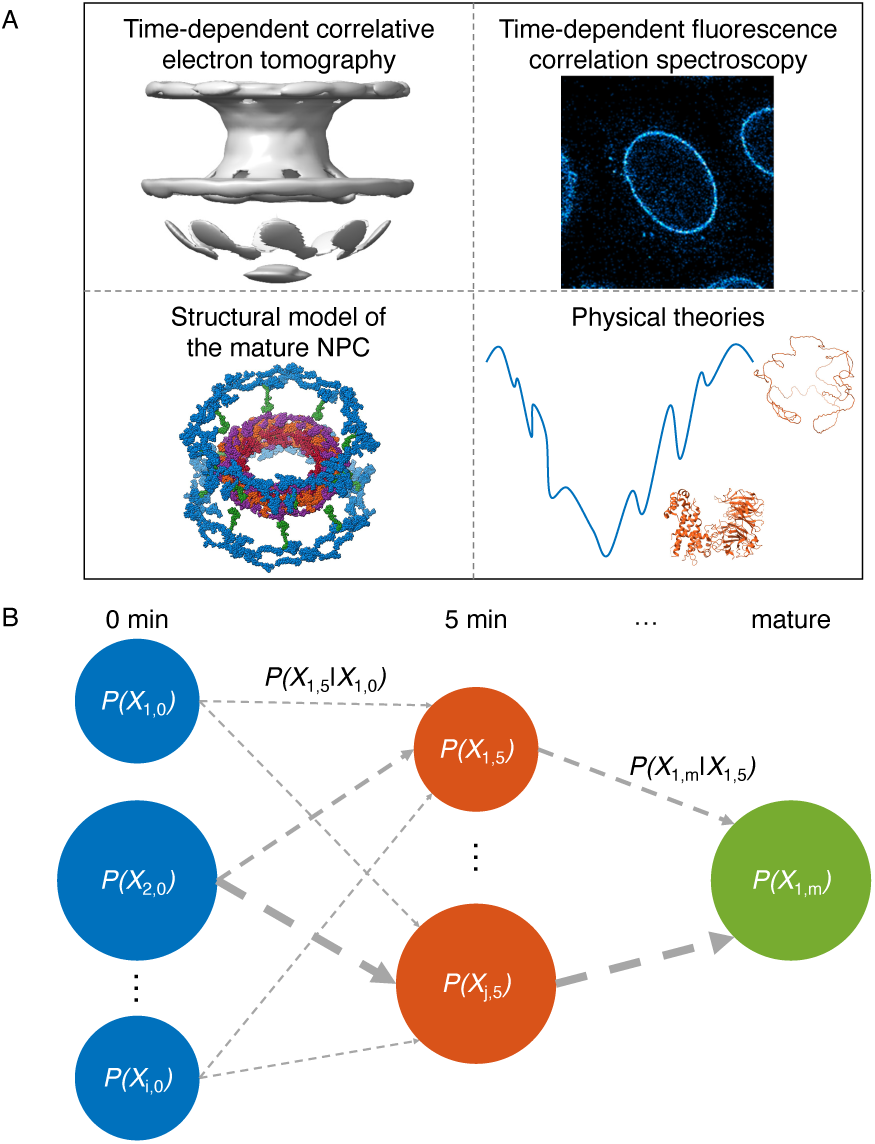
Illustration of integrative spatiotemporal modeling. A) Snapshot models of the NPC intermediates are produced based on structural models of the mature NPC, correlative ET, FCS, and physical theories. B) Trajectory models are created by selecting and connecting snapshots at adjacent time points. The integrative spatiotemporal modeling procedure produces a weighted set of these trajectory models. The weights of both individual snapshots and transitions between these snapshots contribute to the trajectory score (Eq. 2). The weights of individual snapshots are represented by the size of the nodes, while the weights of the transitions are represented by the width of the arrows.

The approach generalizes our method for integrative modeling of static structures, (SI Appendix) including gathering information, defining the model representation, scoring of the models, sampling good scoring models, and validating the models. ^11–13^ Next, we outline each one of these stages in turn, followed by a description of the resulting NPC assembly model.

### Gathering information

In the first stage, available information about the dynamic process of interest is gathered. This information is subsequently used for representing a spatiotemporal model, scoring alternative models, sampling alternative models, filtering sampled models, and/or validating the output models.

For the NPC assembly, information included a model of the mature NPC structure,^45,46^ live-cell correlated electron tomography (ET) maps,^38^ and fluorescence correlation spectroscopy (FCS)-calibrated quantitative fluorescence microscopy data^37^ (SI Appendix, Table S2). The mature NPC structure informed Nup positioning within snapshot models, the scoring of transitions between snapshots, Nup stoichiometry in the final snapshot, and the rigid representation of the NPC subcomplexes. The live-cell correlated ET maps taken at successive time points along the assembly pathway informed the overall size and shape of the assembling NPC as well as the size and shape of the nuclear envelope as a function of time. The quantitative fluorescence microscopy data informed the Nup stoichiometry as a function of time. The collection of all available data is *D* = *{D*_0min_*, …, D*_MP_*}*, where each pair *D_i_*, *D_j_* is mutually independent for *i ̸*= *j*, except for the mature NPC structure. Each snapshot model was scored using the data for that snapshot, *D_i_*.

### Representation

In the second stage, the model representation corresponding to the set of all variables that describe the spatiotemporal dynamics of the system is defined. These variables include spatial coordinates for the components of the system as a function of time. They might also include identifying key snapshot states, molecular composition of a system as a function time, and other “auxiliary” variables, such as data noise parameters. In principle, a wide range of spatial and temporal scales can be represented.

For the NPC assembly, we represented the process by the variables that define a series of snapshot models. Each snapshot model representation included the copy numbers of the Nups in the snapshot (*N_t_*) and their spatial arrangement (*X_N,t_*). We elected to explicitly model only the time points with ET maps: 5 min, 6 min, 8 min, 10 min, 15 min after anaphase onset, and the mature pore structure (SI Appendix, Representation of static snapshots). For completeness, we also show a hypothesis of what the pore may look like at 0 min, which is a pore with the same height and width of what we observe at 5 min, with no Nups present. This hypothesis is uncertain,^37,44^ but does not affect the model of the assembly process from 5 min to the mature pore. In the assembly model, a trajectory is an ordered sequence of snapshot models, *X* = *{*(*X*_0min_*, N*_0min_), …, (*X*_15min_*, N*_15min_), (*X*_MP_*, N*_MP_)*}*. The final set of models is the collection of weighted trajectories, where each trajectory is scored based on the input information (SI Appendix, Table S2).

### Scoring snapshot models

In the third stage, we created a scoring function to weigh candidate trajectory models. To score the structure and composition of static snapshots, we computed and scored snapshot models independently at each time point, using standard integrative modeling of static structures (SI Appendix, Scoring static snapshots and Table S2):

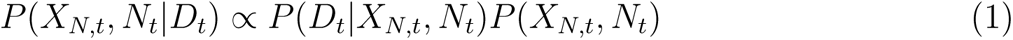

where *P* (*D_t_|X_N,t_, N_t_*) is the data likelihood for the structure and composition of a snapshot and *P* (*X_N,t_, N_t_*) is the prior probability for the structure and composition of a snapshot. These likelihoods and priors reflected the ET maps, FCS data, Gō-like terms,^47^ and excluded volume.

### Scoring trajectory models

Scoring only individual snapshot models along a trajectory generally cannot utilize all information about the process; in particular, it cannot reflect any information about interconversions between different snapshot models. Thus, a complete trajectory model score for a Markovian process^48^ is a product of model scores for each snapshot and each transition in a trajectory (Fig. 1B):

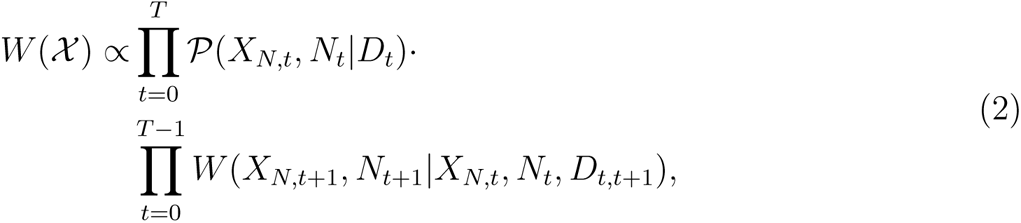

where *t* indexes times from 0 until the final modeled snapshot (*T*); *P*(*X_N,t_, N_t_|D_t_*) is the snapshot model score, which is computed for a constant compositional state and subcomplex location assignment and averages the score of all sufficiently good scoring structural states (Eq. 1); and *W* (*X_N,t_*_+1_*, N_t_*_+1_*|X_N,t_, N_t_, D_t,t_*_+1_) is the transition score. In general, transition scores can be derived from multiple sources of information, including experimental information or physical models of the macromolecular dynamics, such as molecular dynamics simulations,^49–51^ Markov state models,^52–54^ or Bayesian metamodeling.^55^

Here, we scored the transitions between snapshots with a simple metric that either allowed or disallowed a transition. A transition was allowed if the subcomplexes present in first snapshot model were included and had the same subcomplex location assignment in the the second snapshot model (SI Appendix, Fig. S2). This metric was based on the assumptions that Nups are unlikely to dissociate from the assembling complex and that large subcomplex rearrangements are unlikely.

### Sampling

In the fourth stage, we aim to find good-scoring trajectory models. To do so, we need to create a set of alternative snapshot models for each time point, followed by connecting them into a trajectory.

To create a set of alternative snapshot models at all time points, we first enumerated 8-fold symmetric subsets of the mature NPC components. Next, at each time point, we applied the following four-step procedure. First, we ranked each enumeration of the possible Nup copy numbers according to its likelihood (SI Appendix, Eq. S1). All enumerations where the copy number of the Nup205 complex in the inner ring was greater than or equal to that of the Nup188 complex in the inner ring were considered. Second, we selected the top 4 scoring Nup copy enumerations. Third, for each one of these enumerations, we also enumerated the subcomplex location assignments for the Y-complex. Other subcomplexes with multiple copy numbers were randomly assigned an order of subcomplex location assignment before modeling. In total, this procedure resulted in 15, 20, 22, 18, 15, and 1 subcomplex location assignments for, respectively, 5 min, 6 min, 8 min, 10 min, and 15 min after anaphase onset and the mature pore. Finally, for each enumerated subcomplex location assignment, we generated one snapshot model using standard integrative modeling of a static structure (SI Appendix, Sampling static snapshots). This modeling produced a set of alternative structural models of the partially assembled NPC at a given time point.

With multiple snapshot models for each time point in hand, we next connected them into trajectory models. Specifically, we enumerated all connections between snapshot models at adjacent time points, followed by scoring these trajectories according to Eq. 2. The resulting set of trajectory models was represented as a directed acyclic graph. In this representation, each node in the graph is a snapshot model, and each edge represents the transition weight between snapshot models. The complexity of the graph varies with the number of time points (6 for the postmitotic NPC assembly model), the number of snapshot models (91 for the postmitotic NPC assembly model), and the number of edges (425 for the postmitotic NPC assembly model, excluding edges with a weight of 0). These numbers determine the total number of enumerated trajectories (5,184 for the postmitotic NPC assembly model).

### Model analysis and validation

In the fifth stage, we analyze and validate the model of NPC assembly. In general, the uncertainty of the output model results from the actual heterogeneity in the samples used to generate the input information, uncertainty in the input information, and imperfection in the modeling method, which in turn reflects uncertain representation, scoring, and sampling. As for static integrative structure models (SI Appendix, Analysis and validation of static snapshots), we validate the assembly model in four ways, as follows.^13^

First, we analyzed the sampling precision (uncertainty) of the NPC assembly model. The only origin of the sampling uncertainty was the sampling uncertainty of the snapshot models, because the potential assembly pathways were enumerated. To assess how uncertainty originating from the stochasticity of the sampling of snapshot models propagated into the temporal domain, we utilized the Manhattan distance to define the temporal precision of a model:

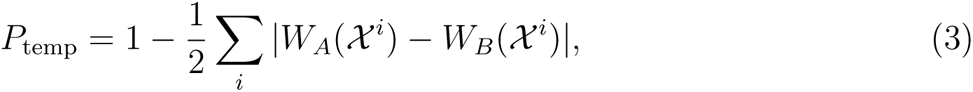

where *i* denotes each possible trajectory (*χ^i^*). Comparisons are made between two independently sampled sets of the same snapshot models (*A* and *B*), each of which has a corresponding weight for a given pathway (*W_A_*(*X ^i^*) and *W_B_*(*X ^i^*), Eq. 2). *P*_temp_ reflects the degree of overlap between different pathway models, taking a value of 1.0 when agreement is perfect and 0.0 when there is no overlap. The metric quantifies the precision (uncertainty) of pathway weights; thus, the weights should only be interpreted up to *P*_temp_.

Second, after assessing the sampling precision of the model, we evaluated the model by comparing it to experimental data used in model construction (SI Appendix, Table S2). In addition to examining the agreement between the snapshot models in the highest weighted trajectory and static data, such as the structure of the assembled NPC and ET maps (SI Appendix, Analysis and validation of static snapshots), we also examined how the set of NPC assembly models matches the FCS data used to score the model. For each Nup used in model scoring (Nup107, Nup93, Nup205, and Nup62), we used the set of trajectory models to compute the weighted sum of Nup copy numbers from each compositional state at each time point. We compared these computed copy numbers to the experimental copy numbers from FCS data.^37^

Third, we evaluated the model by comparing it to experimental data not used in model construction. Specifically, we intentionally excluded Nup copy number data for Nup188 and Seh1 from model construction. Using the same procedure as described in the previous paragraph, we assessed the set of models by comparing the computed copy numbers to the experimental measurements.

Finally, we evaluated the precision (uncertainty) of the set of postmitotic NPC assembly models. The precision of the model is defined by the variation among good-scoring trajectory models:

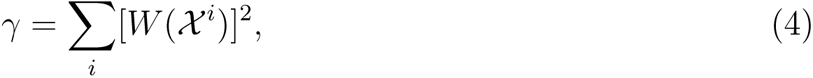

where *i* denotes each possible trajectory (*χ^i^*), each of which has a corresponding weight (*W* (*X ^i^*), Eq. 2). *γ* ranges from 1.0 to 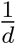, where *d* is the number of trajectories. Values approaching 1.0 indicate that the NPC assembly model can be described by a single trajectory, while values approaching 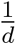 indicate that all trajectories are weighted approximately equally.

## Results

### Molecular model of the postmitotic NPC assembly process

The modeling began by computing snapshot models for the most likely Nup combinations at each sampled time point along the assembly process. A single trajectory model consists of a series of snapshot models, with one snapshot model at each time point. These trajectory models can then be scored by multiplying the scores of snapshot models along the trajectory and the scores of transitions between adjacent snapshots. Thus, the procedure produced a set of weighted trajectory models, with the weights summing to one; the larger the weight of a model, the more consistent the model was with the input information.

The set contained a variety of possible trajectories (SI Appendix, Fig. S3); however, the high model precision (*γ*, Eq. 4) of 0.999 indicated that the set of alternative models can be represented by the single best-scoring trajectory (Fig. 2). This best-scoring trajectory (the pathway model) is the focus of our analysis as follows.

**Figure 2:**
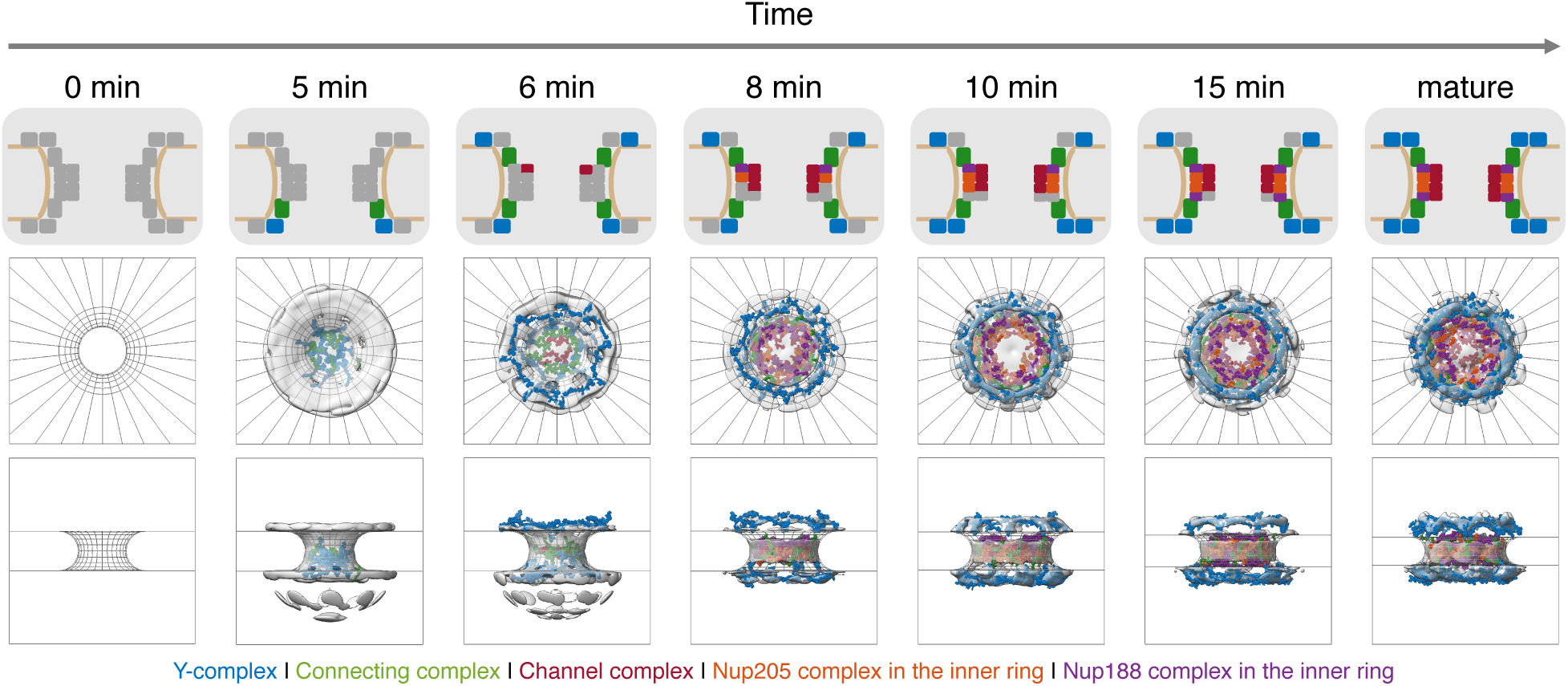
Spatiotemporal model of the postmitotic NPC assembly pathway. The top trajectory, which accounts for 99.9% of model weight, is shown. For each step along the assembly trajectory, a diagram of Nup participation (top panel) is shown, along with a structural model from the cytoplasm (middle panel) and from within the nuclear envelope (bottom panel). Structural models correspond to the centroid structure of the most populated cluster. ET maps are shown in grey, and are compared to structure models corresponding to the Y-complex (blue), connecting complex (green), channel complex (red), Nup205 complex in the inner ring (orange), and Nup188 complex in the inner ring (purple). No experimental data is available at 0 min, corresponding to the completely disassembled NPC in the context of the unsealed nuclear envelope.

In the pathway model, eight Y-complexes initiate assembly on the nuclear side of the nuclear envelope (5 min). Next, the FG-Nup-containing channel complex (Nup62-Nup58-Nup54) localizes toward the center of the pore, as the first cytoplasmic Y-complex is added (6 min). Two more copies of the channel complex (Nup62-Nup58-Nup54) are then added, along with the first components of the inner ring (Nup205-Nup93-Nup155 and Nup93-Nup188-Nup155, 8 min). As assembly proceeds, the inner ring continues to grow, the pore continues to dilate, and the second Y-complex is added to the nuclear side (10 min and 15 min). Eventually, the final copies of the channel complex and cytoplasmic Y-complex are added, leading to the complete NPC structure (mature).

### Assessment of the postmitotic NPC assembly model

A model needs to be assessed before it is interpreted. While the accuracy of a model (the difference between the model and the truth) is often unknown, we can still assess it in at least four ways: estimating the sampling precision, comparing the model to data used to construct it, validating the model against data not used to construct it, and quantifying the precision of the model. These four assessments titrate our confidence in the model; for example, the model precision may be used as a proxy for model accuracy, assuming no systematic errors. Here, each of these assessment steps is exemplified for the postmitotic NPC assembly model. To estimate the sampling precision of the NPC assembly process, we quantified the sampling precision of the only stochastic sampling step, namely that of computing a static snapshot model, and how that sampling precision impacted the uncertainty of trajectory model weights. To estimate the sampling precision of the assembly model, we considered the sampling precisions of the individual snapshots. ^56^ More specifically, we confirmed that two independently obtained samples of each snapshot model had scores drawn from the same parent distribution (Fig. 3A and SI Appendix, Fig. S4) and that each structural cluster from two independently obtained samples included models from each sample proportionally to its size (Fig. 3B and SI Appendix, Fig. S5). We also found *P*_temp_ (Eq. 3) to be 0.9995, indicating that two independent samplings of the models give nearly identical weights to each trajectory.

**Figure 3:**
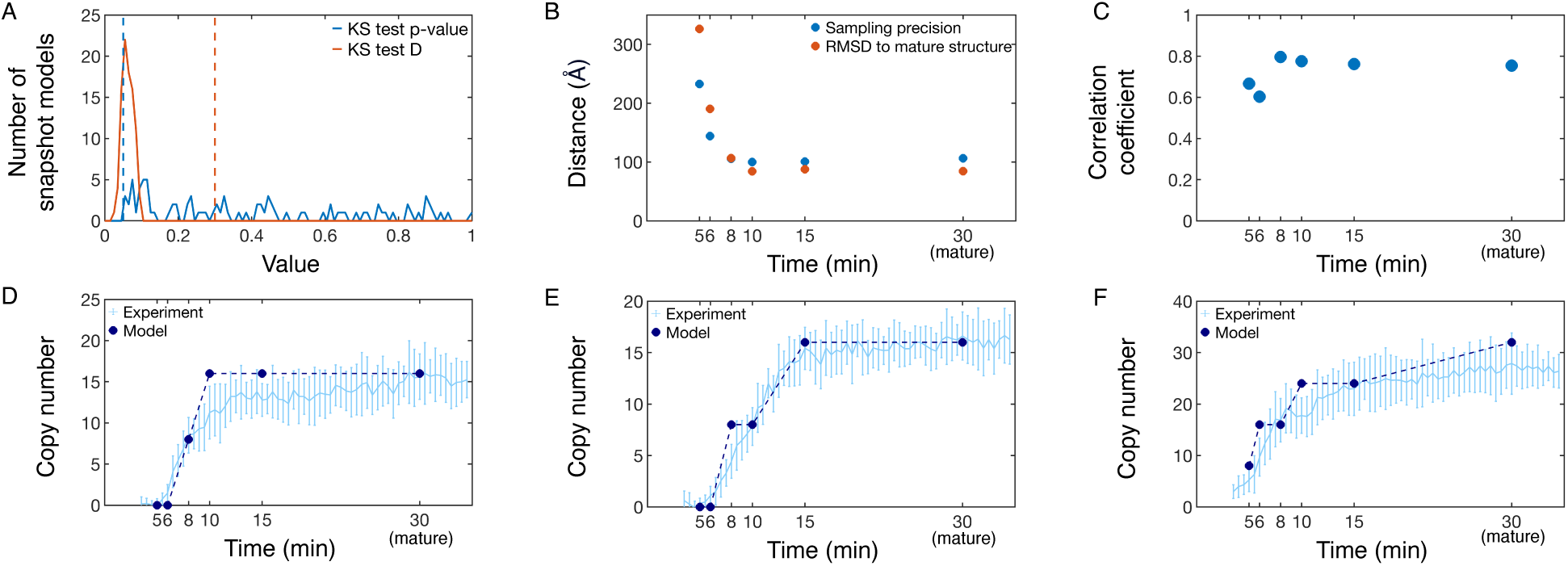
Validation of the spatiotemporal NPC assembly model. A) Determining sampling exhaustiveness of snapshot models along the NPC assembly pathway. For each snapshot model, two samplings are performed and compared via a Kolmogorov-Smirnov two-sample test.^57^ Sampling is performed until the difference in distribution of scores is not significant (p-value *>* 0.05) and small in magnitude (Kolmogorov-Smirnov statistic, D, *<* 0.3). Dashed lines correspond to the cutoffs for Kolmogorov-Smirnov test p-values and Kolmogorov-Smirnov test D. B) Distance-based measures for snapshot models in the best scoring trajectory. Both the structural precision and the average RMSD to the fully mature NPC are shown. C) Correlation coefficient between the snapshot model and the ET map for each snapshot model in the pathway model. D) Comparison of Nup copy number between FCS data (light blue^37^) and the set of NPC assembly trajectory models (dark blue) for Nup205, which is explicitly included in the scoring function. As the FCS curve plateaus by 30 min, we compare the experimental copy number at 30 min to the mature copy number predicted by the model. Error bars represent standard deviations over multiple experimental measurements or over weighted trajectories, and are smaller than the symbols when not visible. The model standard deviation is undefined for the mature state, which we hold fixed. Comparison to other proteins included in the scoring function are found in SI Appendix Fig. S7. Comparisons of Nup copy number between FCS data (light blue^37^) and the set of NPC assembly trajectory models (dark blue) for proteins not explicitly included in the scoring function, Nup188 (E) and Seh1 (F).

To compare the set of trajectory models to data used to construct it, we measured the models’ agreement to the mature structure, the ET maps, and the FCS data. We began by comparing snapshot models at each time point along the pathway model to the mature structure (Fig. 3B). The root mean square deviation (RMSD) to the mature structure followed the same trend as the sampling precision; snapshots at early time points (5 min and 6 min) had a higher uncertainty and deviated slightly from the mature structure, while snapshots at later time points (8 min onwards) were at comparatively higher precision and more closely resembled the mature structure. Next, we calculated the correlation between the snapshot models in the pathway model to the ET data used in model construction (Fig. 3C). We saw good agreement between the snapshot models and the ET maps, with a correlation coefficient of at least 0.60 at all time points. Then, we compared the Nup copy numbers among our weighted sets of trajectory models to experimental copy numbers according to FCS data (Fig. 3D, SI Appendix Fig. S7). While reasonable agreement exists for all Nups, the weakest agreement was observed for Nup93 and Nup62, particularly at later time points. These two Nups were constrained to maximum copy numbers observed in the mature structure,^45,46^ which did not fully account for all copies observed via FCS. This discrepancy demonstrates how our approach is limited by the accuracy of the input information, in this case the mature NPC structure.

To validate the set of trajectory models against the data not used to construct it, we confirmed that our model is consistent with the time dependence of the Nup188 and Seh1 copy numbers determined by FCS (Fig. 3E-F). This test illustrates the predictive power of the model, although the copy numbers of other members of the same subcomplexes were included in model construction.

To quantify the precision of the model, we computed the precision, *γ*, (0.999) and the weight for the most likely trajectory given the data, *W* (*χ*^1^), (99.9%). The high sampling precision of the model ensures that these estimates were not affected by stochasticity of sampling. The high precision indicates that the set of alternative models can be adequately represented by the single, best-scoring trajectory.

### Temporal scoring improves model precision

In principle, integrative modeling can improve the accuracy, precision, and completeness of a model by incorporating more information about a process. To illustrate this point concretely, we quantified the improvement in the precision of the trajectory set afforded by temporal, in addition to spatial, scoring. The trajectory set without temporal scoring is larger because it contains transitions between all pairs of neighboring snapshot models, instead of only those that are allowed in the original set of models (SI Appendix, Fig. S2).

The precision of a trajectory set is visualized via its directed acyclic graph (Fig. 4A). These visualizations clearly indicated the greatly simplified trajectory space for the temporally constrained set compared to the temporally unconstrained set, reducing the number of possible trajectories from 1,782,000 to 5,184. Correspondingly, the unconstrained set was less precise, with a *γ* of 0.36, instead of 0.999 for the constrained set. The difference in precision of the two trajectory sets was also reflected in the weights of the top trajectory (49% and 99.9% for the unconstrained set and constrained set, respectively) and the precision of the Nup copy numbers as a function of time (SI Appendix, Fig. S8).

**Figure 4:**
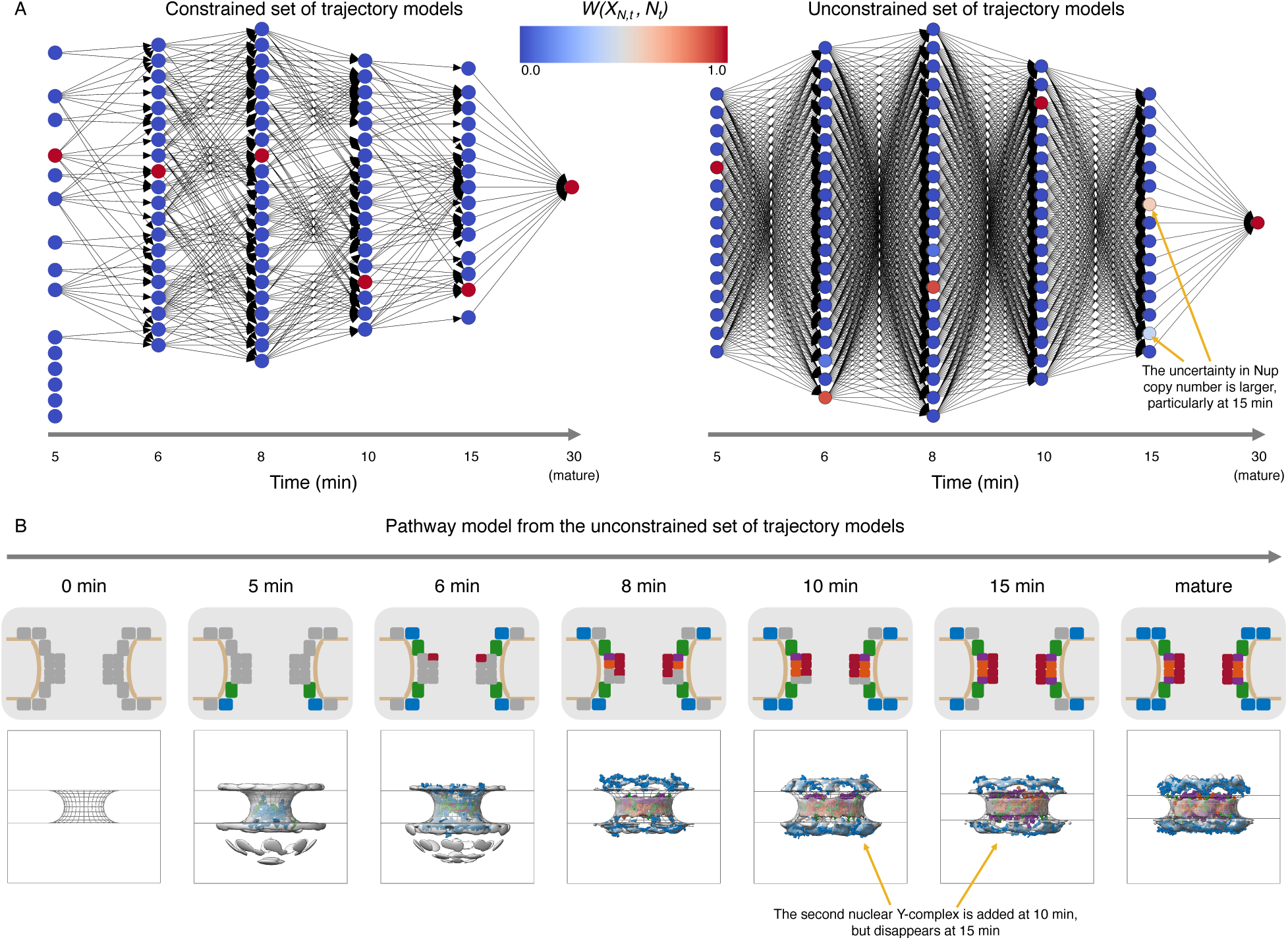
Temporal scoring terms help improve the precision of the resulting model. A) Directed acyclic graphs of the constrained (left) and unconstrained (right) sets of trajectory models. Each column in the graph corresponds to a different time point in the assembly process (5 min, 6 min, 8 min, 10 min, 15 min, and mature). Each node is shaded according to its weight in the final model (*W* (*X_N,t_, N_t_*)). B) Most likely assembly trajectory from the unconstrained set of trajectory models. For each step along the assembly trajectory, a diagram of Nup participation (top panel) is shown, along with a structural model from within the nuclear envelope (bottom panel). Experimental electron density maps are shown in grey, and are compared to protein models corresponding to the Y-complex (blue), connecting complex (green), channel complex (red), Nup205 complex in the inner ring (orange), and Nup188 complex in the inner ring (purple).

Next, we assessed the highest weighted trajectory from unconstrained set of trajectories by comparing it to the highest weighted trajectory from the constrained set of trajectories. As expected, the highest weighted trajectory in the unconstrained set included disallowed transitions in the constrained model. In particular, the second copy of the nuclear Y-complex joined the assembling NPC at 10 minutes, but was then removed at 15 minutes. This rearrangement seems unlikely, further demonstrating how the spatiotemporal constraints term improves the constrained set of models.

## Discussion

We developed integrative spatiotemporal modeling and demonstrated it by modeling the NPC assembly pathway during mitosis. A key advantage of an integrative approach is that it improves the accuracy, precision, and completeness of the model, because it aims to maximize the amount of input information. This point is illustrated by comparing previous schematic models of NPC assembly with our formal, molecular model of NPC assembly: Previous studies probed the assembly of the NPC with individual techniques,^37,38,40^ resulting in a coarse schematic of the composition and shape of the assembling NPC as a function of time.^39^ In contrast, our integrative approach systematically explored “all” Nup trajectories from separate components to the fully assembled mature pore, producing a series of snapshots with defined molecular architectures that were consistent with all input information.

Incorporating additional types of input information into model construction remains a key challenge in spatiotemporal modeling; using information for model construction, as opposed to only filtering or validation, is more likely to result in models consistent with it. In general, input information can come from experiments, physical theories, statistical analyses, and other prior models: for example, diffusion coefficients,^58^ physics-based force fields,^59,60^ meta-modeling,^55^ previous integrative^61,62^ or machine-learned^63–65^ structures of components, protein degradation experiments,^66^ binding rate constants,^67^ molecular networks,^68^ fluorescence microscopies,^69,70^ and electron microscopy.^71,72^

Accordingly, the precision and our confidence in the model of the NPC assembly process could be increased by incorporating additional information into the current method, as illustrated by the following four examples. First, targeted Nups can be depleted through auxin-inducible degradation, ^66^ allowing us to either validate or improve the model based on this information. Specifically, stalled intermediate states would simplify high resolution structural characterization, which could either confirm predictions from the original model or inform new snapshot models.

Second, the completeness of snapshot models could be improved by increasing the resolution of the model representation. For instance, deep-learning models^63–65^ could predict atomistic structures of Nups and their binding modes; furthermore, the nuclear envelope could be represented by a physics based membrane force field,^60^ instead of the current implicit representation.

Third, the precision of the trajectories could be improved by reflecting additional information in the scoring of dynamic transitions between snapshots. For example, such information may be provided by molecular dynamics simulations ^49,50^ or Markov-state modeling.^52–54^

Finally, the completeness of modeling the assembly process of the NPC can be improved by expanding comparative modeling of static structures ^73^ to dynamic processes. In particular, structural models of the mature NPC from multiple organisms ^23–32^ could be integrated with our model of the NPC assembly pathway to compute assembly pathways across different organisms.^55^ Such an analysis may be particularly informative about the evolution of the NPC assembly process and thus the evolution of eukaryotes.^39^

In addition to improving the current method, other spatiotemporal integrative modeling approaches should be developed. In particular, incorporating experimental measurements into Brownian dynamics simulations could in principle result in a set of trajectories that are by design consistent with the input information; the input into such simulations is a starting configuration for a set of components, their interactions, and their diffusion constants. The resulting models would provide a near-continuous and more realistic description of a dynamic process, compared to the current pathways through a series of discrete snapshots. This modeling process could begin by computing Brownian dynamics trajectories that incorporate physics-based diffusion constants^58^ for each molecule, pairwise binding rate constants^67^ between each pair of molecules, and binding modes between each pair of molecules derived from comparative,^73^ integrative,^11–13^ or deep-learning^63–65^ methods. Some temporal experimental information could be incorporated by an iterative search for interaction parameters that produce simulated trajectories whose computed properties match the observed ones. ^74^ Other temporal experimental information may be better incorporated by guiding possible trajectories through enhanced sampling^75–78^ methods. Finally, time-averaged experimental information could be incorporated through biasing potentials such as maximum entropy^79–83^ approaches.

Our model of the NPC assembly pathway provides several insights. Remarkably, modeling shows that one specific trajectory explains the data significantly better than many alternative trajectories. This result is indicative of a specific assembly pathway as opposed to a more stochastic process with multiple, similarly weighted high scoring trajectories. Additionally, as discussed previously,^37^ the model suggests a specific order of assembly for the four copies of the Y-complex per each spoke: the first copy appears on the nuclear side, followed by the second copy on the cytoplasmic side, the third copy on the nuclear side, and finally the fourth copy on the cytoplasmic side. Further, the model suggests that FG-Nups in the central channel complex join the assembling NPC before structured inner ring components. This suggestion, combined with the observation that assembly is initiated by FG-Nups Pom121 and Nup153, leads us to hypothesize that the hydrophobic properties of FG-Nups may be important for the dilation of the nuclear pore and the recruitment of other NPC components.

The spatiotemporal integrative modeling approach suggested here was implemented in our open sourced *Integrative Modeling Platform* (IMP) package,^84,85^ in an effort to facilitate its application by scientists to a wide range of complex biological phenomena.

## Materials and Methods

The spatiotemporal modeling method is outlined in the section entitled “Integrative spatiotemporal modeling workflow”, with additional details in SI Appendix.

## Data Availability

The spatiotemporal modeling method is implemented in our freely available, open source program *Integrative Modeling Platform* (IMP; https://integrativemodeling.org).^84,85^ The input and output files for modeling of the NPC assembly process are available at https://salilab.org/PM NPC Assembly. The NPC assembly model is also available in the PDB archive for integrative structures (XXX). Likewise, the tomograms are also available from the Electron Microscopy Public Image Archive (EMPIAR; EMD-3820) and the FCS date are available at Image Data Resource (IDR; idr0115).

## Acknowledgements

We thank Ben Webb and Ignacia Echeverria for help with implementing our code in IMP. This work was supported by grants from the Baden Wuerttemberg Foundation (J.E. and A.S.), NIH/NIGMS R01GM083960 (A.S.), NIH/NIGMS P41GM109824 (A.S.), NIH/NIGMS R01GM112108 (A.S.) and the European Molecular Biology Laboratory (EMBL; S.O., and J.E.). A.P.L. was further supported by NIH/NIGMS F32GM150243. S.O. was further supported by the EMBL Interdisciplinary Postdoc Programme (EIPOD) under Marie Curie Actions COFUND. S.O. was additionally supported by a Japan Society for the Promotion of Science fellowship (postdoctoral fellowship for research abroad).

