## Supplementary material for "Integrative spatiotemporal modeling of biomolecular processes: application to the assembly of the Nuclear Pore Complex": SI Appendix

### Methods

#### Representation of static snapshots

To represent the structure of the NPC in each snapshot model, we divided the partial mature NPC structure (PDB: 5a9q, 5ijn) into a set of rigid subcomplexes (SI Appendix, Table S3).[S1,S2] Each subcomplex was coarse-grained from the modeled structure of the mature NPC (PDB: 5a9q, 5ijn) by grouping 10 consecutive residues into one coarse-grained bead. The copy number of the subcomplexes in each snapshot is described by $N_t$, which is a vector of integer values specifying the copy number of each subcomplex present at time $t$. The copy number of the connecting complex with Nup155, for which no copy number data is available, was coupled to the first appearing Y-complex on the same side of the nuclear pore. The spatial configuration of the NPC subcomplexes is the vector of Euclidean coordinates, $X_{N,t}$, given the copy number, $N_t$. The nuclear envelope was represented as a fixed toroidal surface embedded in two parallel planes. We set the inner pore diameter and minor radius of the nuclear envelope at each time point to the mean of previously determined nuclear envelope cross sections[S3] with a pore diameter of 51.53 nm, 58.39 nm, 72.74 nm, 84.59 nm, 79.83 nm, and 87 nm; and minor radius of 21.36 nm, 21.245 nm, 21.495 nm, 20.27 nm, 17.125 nm, and 15 nm for the time points at 5 min, 6 min, 8 min, 10 min, 15 min, and the mature pore, respectively. Therefore, the joint set of variables, $(X_{N,t}, N_t)$, fully describes the compositional and structural state of the assembling NPC.

#### Scoring static snapshots

In this section, we detail the likelihood and prior terms that are used to score static snapshot models.

We defined the data likelihood of the compositional state, $P(D_t|N_t)$, as

$$P(D_t|N_t) \propto \prod_{l} \frac{1}{\sigma_l\sqrt{2\pi}} e^{-\frac{(n_{t,l}-\mu_l)^2}{2\sigma_l^2}} \tag{S1}$$

where $\mu_l$ and $\sigma_l$ are the mean and variance of the average copy number of fluorescently tagged Nups estimated in a previous study[S4] and $l$ indexes the list of tagged Nups that are included in the representation $l \in \{Nup,\}$. We used 4 Nups directly in the spatiotemporal scoring (Nup107, Nup93, Nup205, and Nup62), and reserved two Nups for model validation (Nup188 and Seh1).

We scored the shape of the assembling NPC by the cross-correlation between the ET density map[S3] and the assembling NPC structural state. We represented the forward model by fitting each Nup subcomplex with a Gaussian mixture model (GMM) of two components per subcomplex using the *gmconvert* utility.[S5] We represented the ET protein densities at each time with a GMM with 150 components fit using the same tool.

We scored each structural state of the assembling NPC by comparing it to the mature NPC structure[S1,S2] through a harmonic Gō-like model.[S6] Each 10-residue coarse-grained bead was restrained to its position in the mature structure, according to

$$-\ln[P(X_{N,t})] \propto U_{\mathrm{G\bar{o}}}^{\mathrm{prot}} = \sum_i \frac{(\vec{x}_i - \vec{x}_i^{\mathrm{MP}})^2}{\sigma}, \tag{S2}$$

where $U^{\mathrm{prot}}$ is the energy of the snapshot configuration, $i$ is each coarse grained bead, $\vec{x}_i$ is the position of that bead in the sampled structure, $\vec{x}_i^{\mathrm{MP}}$ is the position of that bead in the mature pore, and $\sigma$ is a nuisance parameter that defines the strength of the interaction. Early in the assembly process (specifically at t=5 min), there is a poor correlation between the assembled NPC structure and the ET data (SI Appendix, Fig. S1). As such, we utilized $\sigma = 10^7$ (kcal/mol)$^{-1}$ Å$^2$ at 5 min, and strengthened the Gō interactions by using $\sigma = 10^6$ (kcal/mol)$^{-1}$ Å$^2$ at all other time points.

To ensure the reference pore positions maximally correlate to the ET data at a given time point, we fit the rigid mature structure to the ET map at each time point, and used the best fitting structure as the reference structure ($\{\vec{x}_i^{\mathrm{MP}}\}$). After centering the structure and ET at the origin and aligning the structure along the Z-axis, we enumerated 0.1 nm translations between 5 and -5 nm, and 1 degree rotations of 1-45 degrees, in accordance with the eight-fold symmetry of the NPC. We expanded the search in translational space to 10 and -10 nm for the 5 min case due to a lack of a local maximum between 5 and -5 nm. For each translation-rotation combination, we compared the forward ET density from the GMM,[S5] to the experimental density map using the cross-correlation coefficient. We then based the restraint in Eq. S2 on the mature structure that best fit the experimental ET density at that time point (SI Appendix, Fig. S1).

We also included other physics based terms, based on knowledge of the NPC and its environment in the nuclear envelope. Excluded volumes between subcomplexes were scored as a harmonic repulsion term between CG beads, and were slowly added throughout the three step sampling process. At the first step, beads had no excluded volume. In the second step, spherical proteins beads were given excluded volume (strength 0.01 kcal/mol Å$^2$). Finally, in the final step, the protein excluded volume was replaced with excluded volume of coarse-grained beads (strength 10 kcal/mol Å$^2$). Contacts between Nup domains containing an ALPS-motif were restrained by a harmonic function (strength 0.001 kcal/mol Å$^2$) to be in contact with the nuclear envelope surface. Overlap between the Nups and the nuclear envelope was scored by imposing a harmonic repulsion between the Nups and nuclear envelope surface (strength of 0.01 kcal/mol Å$^2$).

Finally, we combined these scoring terms to compute the Bayesian posterior of a snapshot model according to Eq. 1. In practice, we recognize that, for NPC assembly snapshot models, the data likelihood ($P(D_t|X_{N,t}, N_t)$) is dependent on the Nup copy number, but not the structural state, while the prior probability ($P(X_{N,t}, N_t)$) is dependent on the structural state, and are computed at a fixed Nup copy number by definition. To simplify the trajectory model, we computed a single snapshot model score for each snapshot as:

$$\mathcal{P}(X_{N,t}, N_t|D_t) \propto P(D_t|N_t)\mathcal{P}(X_{N,t}) \quad \text{(S3)}$$

where $P(D_t|N_t)$ are the model likelihoods, which are dependent on Nup copy number and are constant across all structural states, and $\mathcal{P}(X_{N,t})$ are the prior probabilities from the averaged energies, which are dependent on the structural states, and are computed at a fixed Nup copy number. $\mathcal{P}(X_{N,t})$ was extracted from the Boltzmann average of the structural scores:

$$\mathcal{P}(X_{N,t}) \propto e^{\frac{-<U(X_{N,t})>}{k_B T}}, \tag{S4}$$

where $< U(X_{N,t}) >$ is the average energy of the snapshot model, $k_B$ is Boltzmann's constant, and $T$ is a nuisance parameter that balances the scaling of the likelihoods and priors. We propose that the geometric mean of the temperatures at which sampling is performed is a reasonable choice for $T$ (671 K for the NPC assembly model).

#### Sampling static snapshots

For each of the identified subcomplex location assignment and time combinations, we initiated at least 200 three-step temperature replica-exchange Markov Chain Monte Carlo simulations. All steps used random translation and rotation of rigid bodies, with moves up to 1 Å and 0.01 radians, respectively. Each step enforced different levels of protein excluded volume, with all other terms held constant. This choice allowed us to relax the other terms in the scoring function before gradually adding in excluded volume.

For the first step, excluded volume between protein beads was ignored at all levels of the hierarchy. The simulation began with Nups distributed randomly throughout the simulation box, used 8 replicas geometrically spaced between 300 and 450 K, and attempted exchanges every 1000 steps. Simulations lasted for $10^6$ steps, and configurations were saved every 1000 steps. The lowest energy conformation across all replicas was then used as the starting point for the second step.

In the second step, excluded volume was enforced at the protein level of the hierarchy. Starting from the lowest energy configuration from the first step, we used 8 replicas geometrically spaced between 300 and 1500 K, and attempted exchanges every 1000 steps. This second step lasted for $2 \times 10^5$ steps, and configurations were saved every 1000 steps. The lowest energy configuration across all replicas was again saved as the starting point for the third step.

Finally, the third and final step used excluded volume at the smallest coarse-grained bead level of the hierarchy, which typically had 10 residues per bead. We started at the lowest energy configuration from the second step, used 8 replicas geometrically spaced between 300 and 1500 K, and attempted exchanges every 1000 steps. This second step lasted for $2 \times 10^5$ steps, and configurations were saved every 1000 steps. The lowest energy configuration from each replica was saved as as a candidate for good scoring configurations.

To find good-scoring configurations for a given snapshot model, we began with a collection of all of the lowest energy configurations from each replica across all independent sampling runs. To remove poorly-scoring models, we found the median score for the lowest energy configurations from all replicas from all replica-exchange Markov Chain Monte Carlo simulations and selected all models whose score was less than or equal to that median score.

In total, we sampled at least 2,240,000 structural states for each snapshot model (231,392,000 total structural states for all snapshot models), with at least 801 good-scoring structural states per snapshot model (82,732 total good-scoring structural states across all snapshot models). All simulations were performed in IMP.[S7,S8]

#### Analysis and validation of static snapshots

For stochastic sampling methods, thoroughness of sampling can be assessed by quantifying the highest precision sufficient for which sampling is exhaustive. To assess the sampling exhaustiveness of each snapshot model, we followed previous tests for integrative modeling of static structures.[S9] We started by dividing each snapshot model into two independently sampled models, and performing a non-parametric Kolmogorov–Smirnov (KS) two-sample test.[S10] For each snapshot model, we used this test to ensure that the difference between the model scores was statistically insignificant (p-value $> 0.05$) and small in magnitude (KS two sample test statistic, D, $< 0.30$), as shown in Fig. 3A. Then, we aimed to quantify the sampling precision for in configuration space for snapshots along the most likely NPC assembly trajectory. To assess the spatial precision, we utilized the one-sided $\chi^2$ test for the homogeneity of proportions between independently sampled structures. For each pair of configurations, we performed threshold-based RMSD clustering at 1Å intervals. Clustering was considered converged when 1) $\chi^2$ p-value $> 0.05$, 2) Cramer's V $< 0.10$, and 3) the clustered population $> 0.80$ (SI Appendix, Fig. S5).

After assessing the sampling precision of the model, we compared the pathway model to data used in model construction (SI Appendix, Table S2). Some restraints such as protein excluded volume, transmembrane restraint, and membrane exclusion restraint, were implemented through harmonic potentials. For these restraints, they were deemed satisfied if they were violated by less than 3 standard deviations, defined as $\sigma = \sqrt{\frac{1}{k}}$, where $k$ is the spring constant of the harmonic potential.

Comparisons between the mature NPC and the model were made by measuring the RMSD between the mature, aligned NPC (SI Appendix, Fig. S1) and each structure in each snapshot model along the pathway model. Only Nups present in the intermediate structure were considered, and the average was taken over all structures in each snapshot (Fig. 3B).

Final comparisons to ET maps were made by predicting the forward density map directly from the particle positions. The correlation between the ET map and the forward model was evaluated for each snapshot model in the pathway model (Fig. 3C).

### Comparison to a previous model of postmitotic NPC assembly

The model presented here differs slightly from the one presented previously (PDB-dev:00000142).[S4] While some changes originate from modifications to the scoring function and sampling procedure to better allow for an estimation of model precision, other differences originate from a error found in the original code for selecting protein copy numbers. Fixing this error allows the model to better reproduce the FCS data (SI Appendix, Fig. S9). Further, this defect in the previous model demonstrates the need to carefully evaluate a model against the input information used to construct it. The current model was also deposited into PDB-dev with a PDB code of XXX. None of the model differences impact the discussion and conclusions in the original publication.[S4]

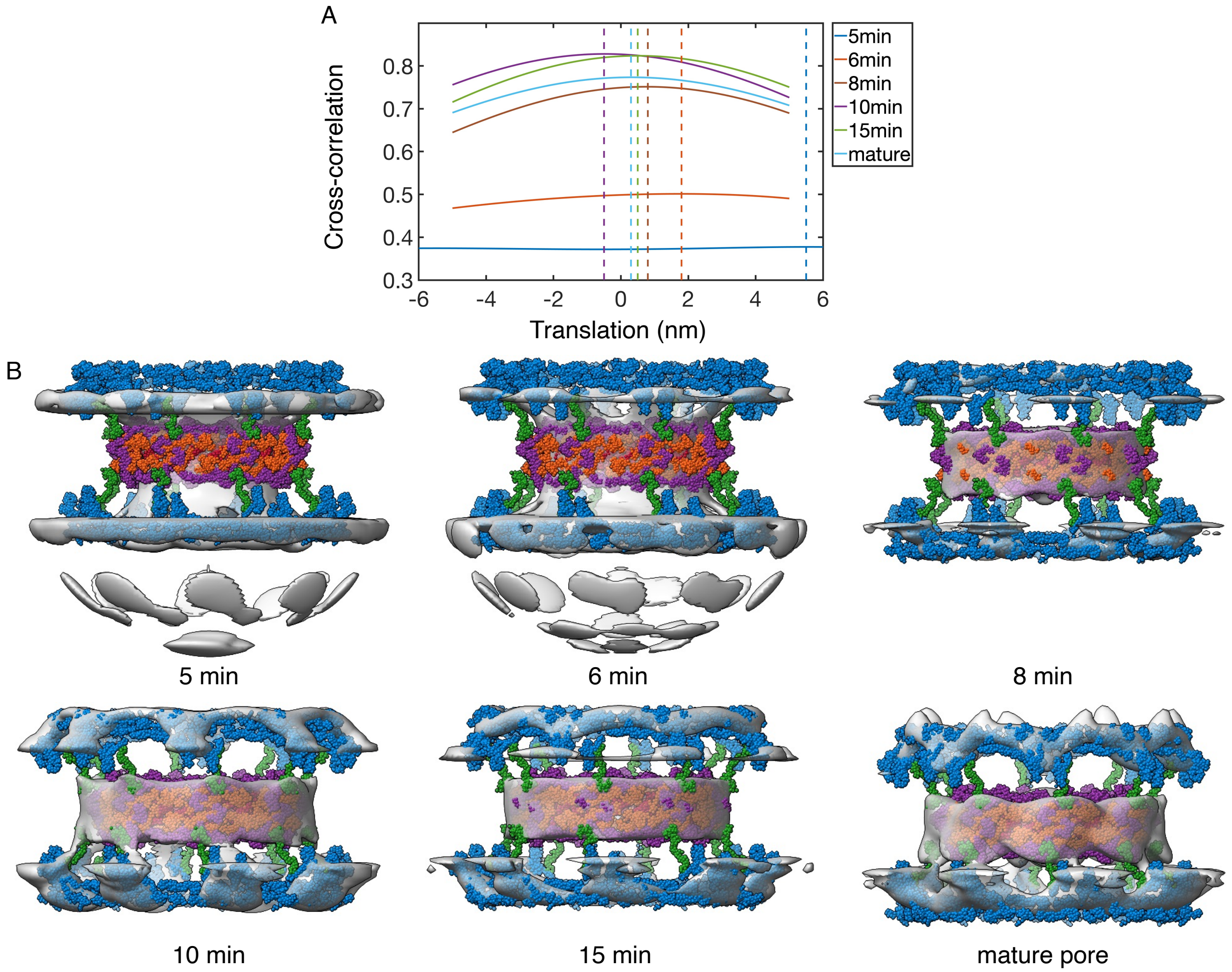


Figure S1: Enumerating fits of the mature NPC structure to time-dependent ET maps. A) Cross-correlation between the forward Gaussian mixture model and that from ET data as a function of translating the mature pore structure along the Z-axis. Values shown are the maximum of 45 possible rotations. Negative values translate the NPC into the nucleoplasm, while positive values translate the NPC into the cytoplasm. Dashed vertical lines represent the translation that maximizes the cross-correlation. B) Fitted structures aligned to the ET map (gray) at each time point. Proteins are colored as Y-complex (blue), connecting complex (green), channel complex (red), Nup205 complex in the inner ring (orange), and Nup188 complex in the inner ring (purple).

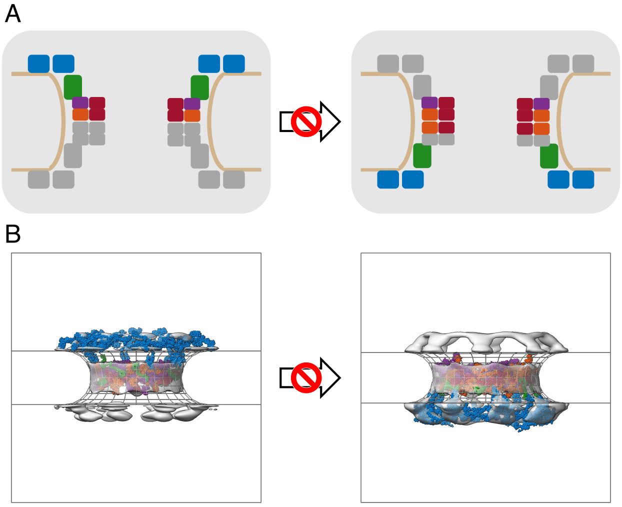


Figure S2: Example of a disallowed transition in the spatiotemporal model. The NPC is represented as both a diagram of Nup participation (A) and a structural model from the cytoplasm (B). Here, this transition is disallowed because the both nuclear Y-complexes are present in the starting state, but not in the ending state. Experimental ET maps (grey), Y-complex (blue), connecting complex (green), channel complex (red), Nup205 complex in the inner ring (orange), and Nup188 complex in the inner ring (purple).

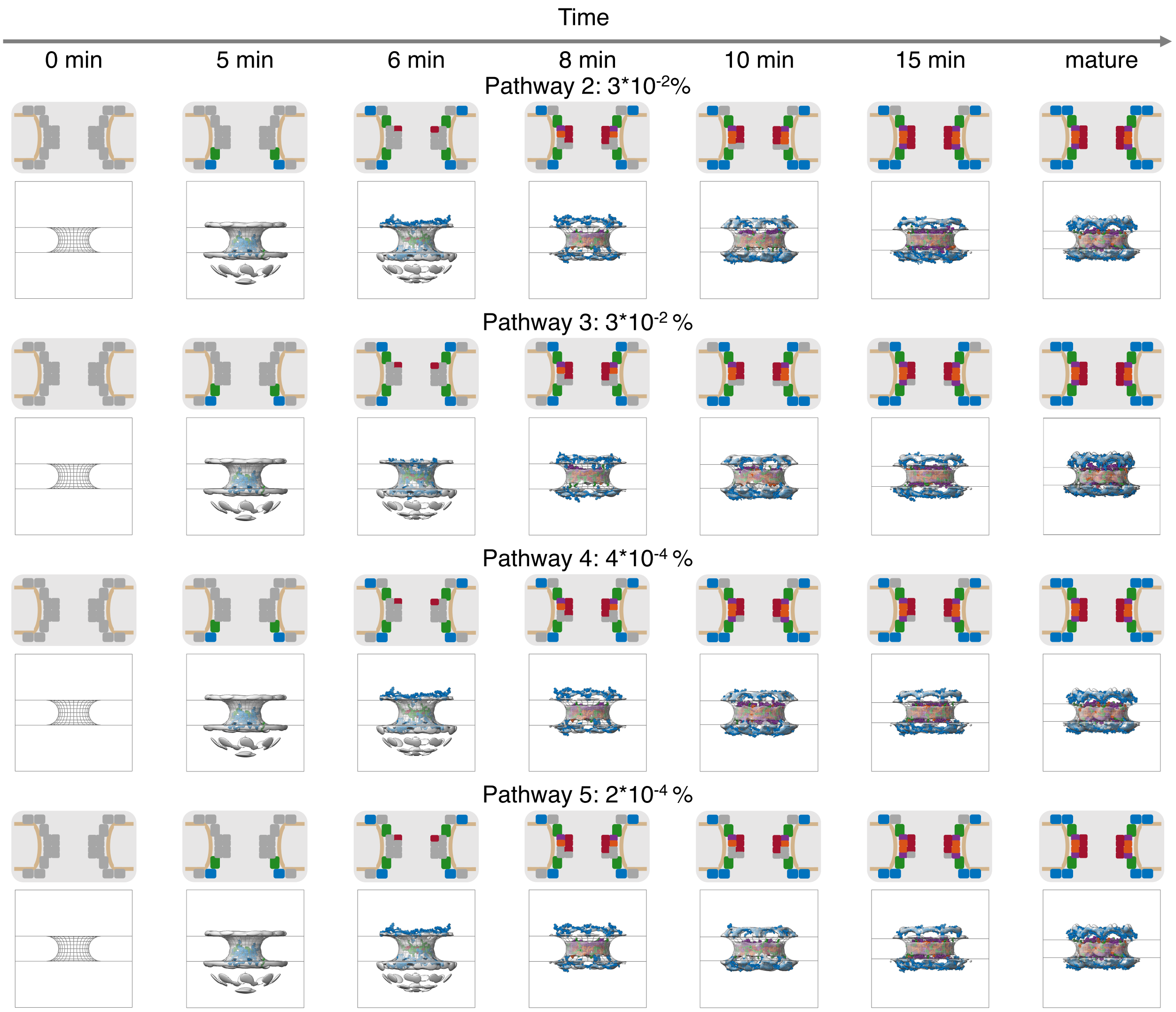


Figure S3: Secondary trajectories in the spatiotemporal model of postmitotic NPC assembly. The second to fifth most likely trajectories are shown, and can be compared to the most likely trajectory, which accounts for 99.9% of the model weight and is shown Figure 2. Each trajectory includes its weight in the final assembly model, stated as a percent of all possible trajectories. For each step along the each trajectory, a diagram of Nup participation (top panel) is shown, along with a structural model from within the nuclear envelope (bottom panel). Structural models correspond to the centroid structure of the most populated cluster. Experimental electron density maps are shown in grey, and are compared to protein models corresponding to the Y-complex (blue), connecting complex (green), channel complex (red), Nup205 complex in the inner ring (orange), and Nup188 complex in the inner ring (purple). No experimental data is available at 0 min, which is not explicitly modeled.

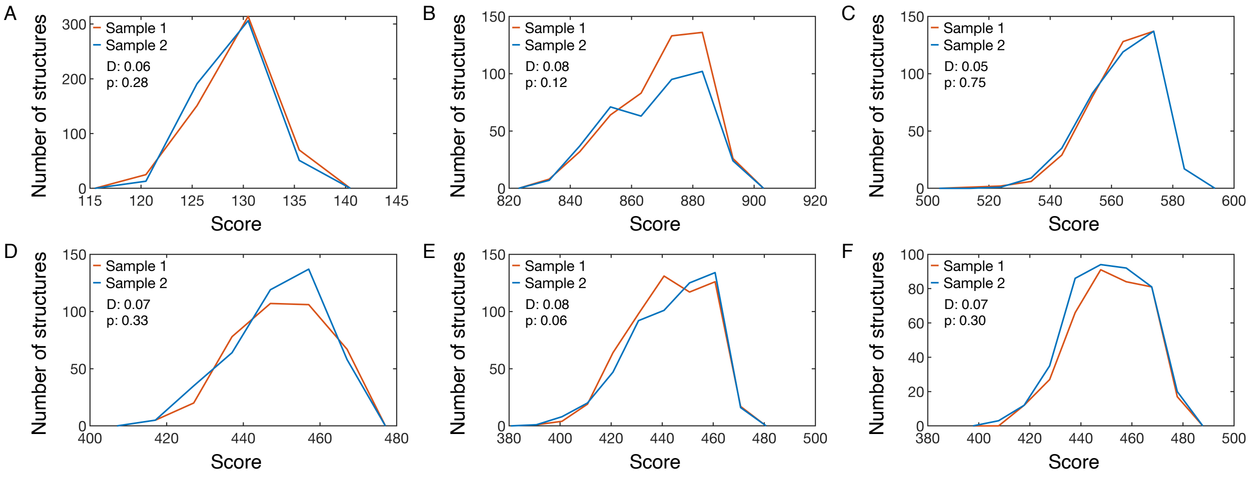


Figure S4: Example of determining sampling exhaustiveness for snapshots along the pathway model. For a given Nup configuration at a given time point, we compare two independent samplings of snapshot models. The difference between the two samplings is measured using a KS two-sample test, and sampling is performed until the difference in distribution of scores is not significant (p-value, p, $> 0.05$) and small in magnitude (KS statistic, D, $< 0.3$).

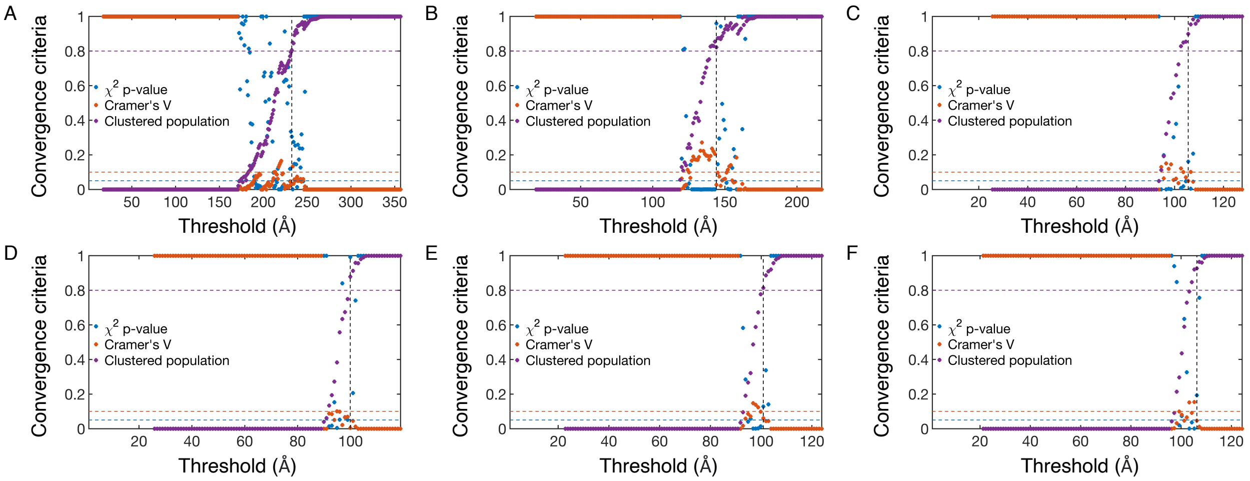


Figure S5: Determination of sampling precision for the pathway model. For a given Nup configuration at a given time point, we utilize threshold based clustering at 1 Å intervals, and utilize the smallest clustering threshold where 1) $\chi^2$ p-value $> 0.05$ (blue), 2) Cramer's V $< 0.10$ (orange), and the clustered population $> 0.80$ (purple).[S9] Clustering is performed at 5 min (A), 6 min (B), 8 min (C), 10 min (D), 15 min (E), and mature (F).

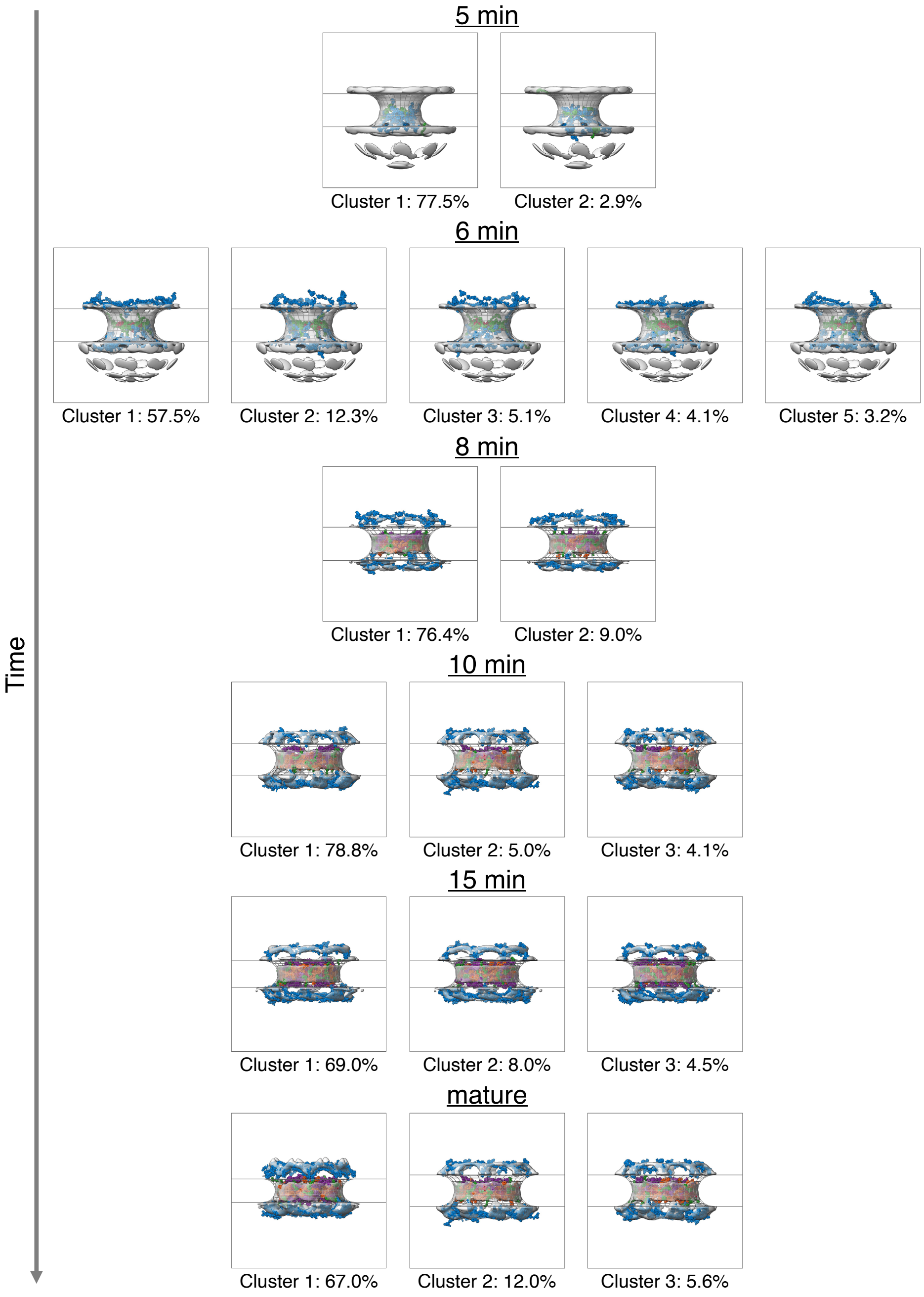


Figure S6: Structural clustering of snapshots along the pathway model. The centroid configuration for each cluster is demonstrated for each step along the assembly process from within the nuclear envelope. Percentage values indicate the population of each cluster from the overall snapshot model. Experimental electron density maps are shown in grey, and are compared to protein models corresponding to the Y-complex (blue), connecting complex (green), channel complex (red), Nup205 complex in the inner ring (orange), and Nup188 complex in the inner ring (purple).

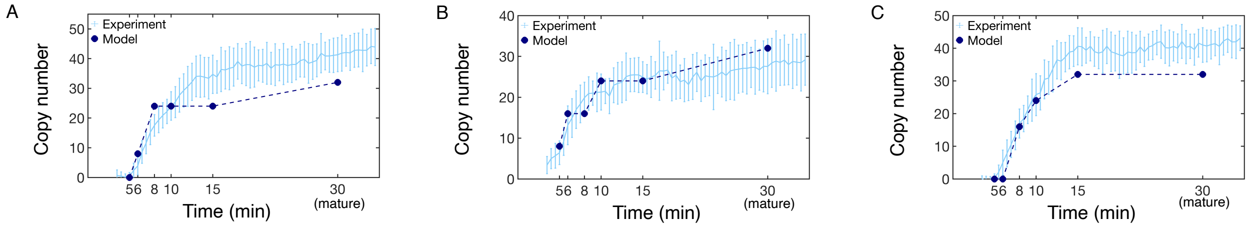


Figure S7: Comparison of Nup copy number between FCS data (light blue[S4]) and the set of NPC assembly trajectory models (dark blue) for Nups explicitly included in the scoring function (A) Nup62, B) Nup107, and C) Nup93). As the FCS curve plateaus by 30 min, we compare the experimental copy number at 30 min to the mature copy number predicted by the model. Error bars represent standard deviations over multiple experimental measurements or over weighted trajectories, and are smaller than the symbols when not visible. The model standard deviation is undefined for the mature state, which we hold fixed.

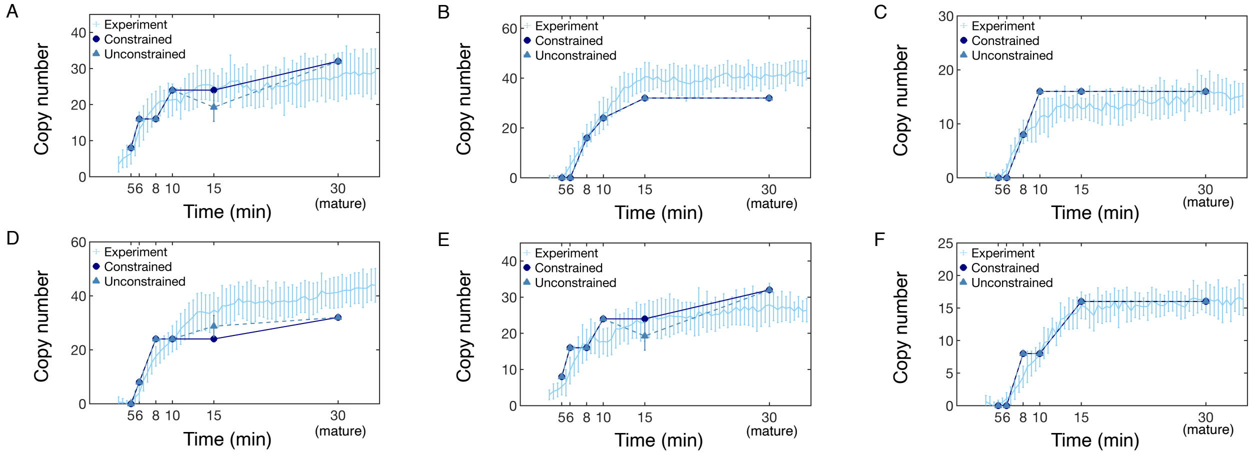


Figure S8: Comparison of Nup copy number between FCS data (light blue[S4]), the constrained set of NPC assembly trajectory models (circles), and the unconstrained set of NPC assembly trajectory models (triangles). Nups explicitly included in the scoring function, A) Nup107, B) Nup 93, C) Nup205, and D) Nup62, and not explicitly included in the scoring function, E) Seh1 and F) Nup188, are shown. As the FCS curve plateaus by 30 min, we compare the experimental copy number at 30 min to the mature copy number predicted by the model. Error bars represent standard deviations over multiple experimental measurements or over weighted trajectories, and are smaller than the symbols when not visible. The model standard deviation is undefined for the mature state, which we hold fixed.

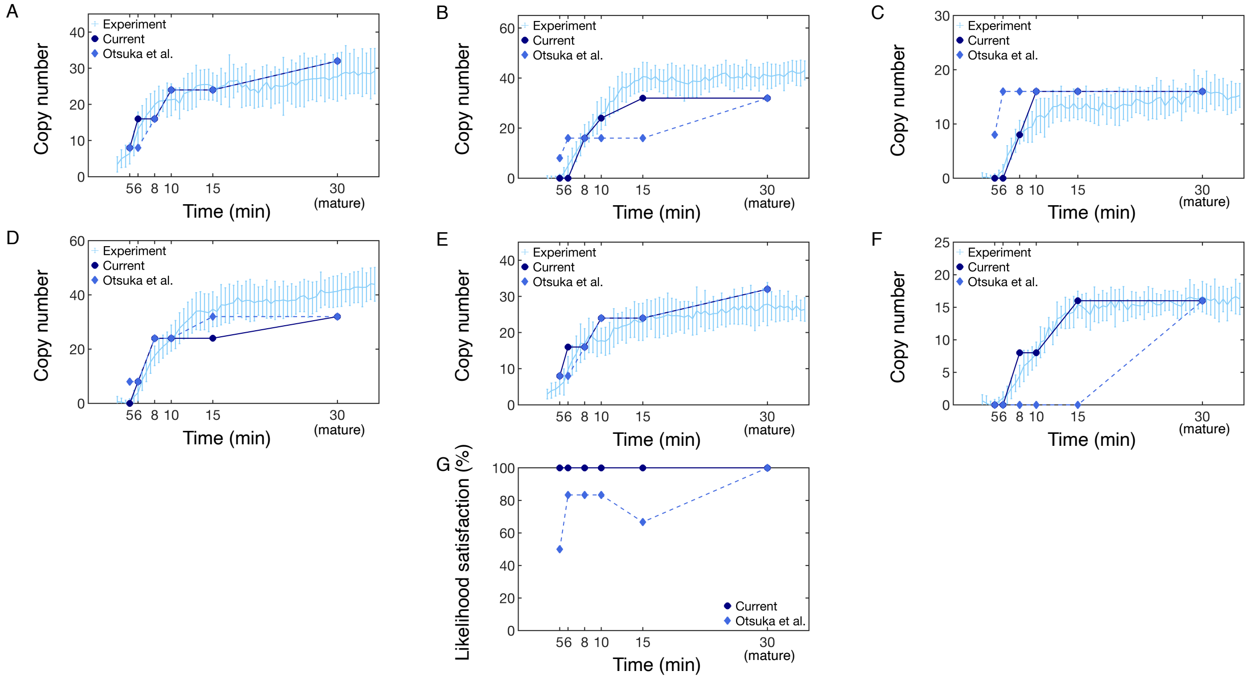


Figure S9: Comparison of Nup copy number between FCS data (light blue[S4]), the set of NPC assembly trajectory models from the current work (circles), and the previous set of NPC assembly trajectory models from Otsuka et al.[S4] (diamond). Nups explicitly included in the scoring function, A) Nup107, B) Nup 93, C) Nup205, and D) Nup62, and not explicitly included in the scoring function, E) Seh1 and F) Nup188, are shown. As the FCS curve plateaus by 30 min, we compare the experimental copy number at 30 min to the mature copy number predicted by the model. Error bars represent standard deviations over multiple experimental measurements or over weighted trajectories, and are smaller than the symbols when not visible. The model standard deviation is undefined for the mature state, which we hold fixed. G) Likelihood satisfaction is defined as the percentage of Nup copy numbers from the set of models that are within 3 standard deviations of the experimental mean. 100% indicates good agreement between the FCS data and the set of models.

Table S1: Summary of Integrative structure determination.

| | |
|---|---|
| Software for modeling the NPC structure | Integrative Modeling Platform (IMP, version 2.20)[S7,S8] |
| Pore membrane representation | Half-torus, whose the diameter (D) and minor radius (r) is based on ET data[S3]<br>5-min D: 51.53 nm r: 21.36 nm<br>6-min D: 58.39 nm r: 21.245 nm<br>8-min D: 72.74 nm r: 21.495 nm<br>10-min D: 84.59 nm r: 20.27 nm<br>15-min D: 79.83 nm r: 17.125 nm<br>mature D: 87 nm r: 15 nm |
| Selection of Nup stoichiometry | Sampled the top 4 scoring Nup combinations for each time point. |
| Configurational sampling method | At least 200 independent Markov Chain Monte Carlo simulations for each snapshot model, starting from random structures.<br>Each independent Markov Chain Monte Carlo simulation consisted of three steps of<br>temperature replica-exchange simulations, where each step used a different representation of excluded volume:<br>Step 1: No excluded volume.<br>$10^6$ steps with 8 replicas between 300 and 450 K, and configurations saved every 1000 steps.<br>Step 2: Excluded volume for each coarse grained protein.<br>$2 \times 10^5$ steps with 8 replicas between 300 and 1500 K, and configurations saved every 1000 steps.<br>Step 3: Excluded volume for each 10 residue coarse grained bead.<br>$2 \times 10^5$ steps with 8 replicas between 300 and 1500 K, and configurations saved every 1000 steps. |
| Monte Carlo moves | Random translation and rotation of rigid bodies, up to 1 Å and 0.01 radians, respectively. |
| Number of modeled structures generated | at least 2,240,000 structural states for each snapshot model<br>(231,392,000 total structural states for all snapshot models)<br>at least 801 good-scoring structural states per snapshot model<br>(82,732 total good-scoring structural states across all snapshot models) |
| Sampling precision | The smallest clustering threshold where independent runs resulted in<br>1) $\chi^2$ p-value $> 0.05$, 2) Cramer's V $< 0.10$, and 3) the clustered population $> 0.80$.[S9]<br>See Fig. S5 and Fig. 3B. |
| Temporal precision | The temporal precision ($P_{\text{temp}}$), defined as the pathway overlap between<br>independent realizations of the model (Eq. 3), was 0.9995. |
| Visualization and plotting | MATLAB and UCSF ChimeraX[S11] |

Table S2: Data representation, scoring, and satisfaction. *: quantities determined by harmonic restraints, with spring constants of 10 kcal/mol Å, 0.001 kcal/mol Å, and 0.01 kcal/mol Å for the protein excluded volume, transmembrane, and membrane exclusion restraints respectively. Satisfaction is defined as being violated by less than 3 standard deviations.

| Method | Scoring | Satisfaction |
|---|---|---|
| Steric effect | Protein excluded volume restraint | 100% satisfied[1] |
| NPC static structure (PDB:5a9q,[S1] 5ijo[S2]) | RMSD restraint, Eq. S2 | Figure 3B |
| | Transmembrane restraint | 100% satisfied[1] |
| ET | Membrane exclusion restraint | 100% satisfied[1] |
| | Shape of the NPC | Figure 3C |
| FCS | Nup stoichiometry likelihood, Eq. S1 | Figure 3D, Figure S7 |

Table S3: Division of rigid subcomplexes used in integrative modeling of the NPC assembly pathway.

| Subcomplex | Protein | PDB:ChainID |
|---|---|---|
| Nuclear Y-complex (inner) | Nup133 | 5a9q:L |
| | Nup107 | 5a9q:M |
| | Nup96 | 5a9q:N |
| | Sec13 | 5a9q:O |
| | Seh1 | 5a9q:P |
| | Nup85 | 5a9q:Q |
| | Nup43 | 5a9q:R |
| | Nup160 | 5a9q:S |
| | Nup37 | 5a9q:T |
| Nuclear Y-complex (outer) | Nup133 | 5a9q:C |
| | Nup107 | 5a9q:D |
| | Nup96 | 5a9q:E |
| | Sec13 | 5a9q:F |
| | Seh1 | 5a9q:G |
| | Nup85 | 5a9q:H |
| | Nup43 | 5a9q:I |
| | Nup160 | 5a9q:J |
| | Nup37 | 5a9q:K |
| Cytoplasmic Y-complex (inner) | Nup133 | 5a9q:3 |
| | Nup107 | 5a9q:4 |
| | Nup96 | 5a9q:5 |
| | Sec13 | 5a9q:6 |
| | Seh1 | 5a9q:7 |
| | Nup85 | 5a9q:8 |
| | Nup43 | 5a9q:9 |
| | Nup160 | 5a9q:a |
| | Nup37 | 5a9q:b |
| Cytoplasmic Y-complex (outer) | Nup133 | 5a9q:U |
| | Nup107 | 5a9q:V |
| | Nup96 | 5a9q:W |
| | Sec13 | 5a9q:X |
| | Seh1 | 5a9q:Y |
| | Nup85 | 5a9q:Z |
| | Nup43 | 5a9q:0 |
| | Nup160 | 5a9q:1 |
| | Nup37 | 5a9q:2 |
| Nup93-Nup205-Nup155 inner ring complex (repeat 1) | Nup93 | 5ijo:C |
| | Nup205 | 5ijo:D |
| | Nup155 | 5ijo:E |
| Nup93-Nup205-Nup155 inner ring complex (repeat 2) | Nup93 | 5ijo:O |
| | Nup205 | 5ijo:P |
| | Nup155 | 5ijo:Q |
| Nup93-Nup188-Nup155 inner ring complex (repeat 1) | Nup93 | 5ijo:I |
| | Nup188 | 5ijo:J |
| | Nup155 | 5ijo:K |
| Nup93-Nup188-Nup155 inner ring complex (repeat 2) | Nup93 | 5ijo:U |
| | Nup188 | 5ijo:V |
| | Nup155 | 5ijo:W |
| Nup62-Nup58-Nup54 channel complex (repeat 1) | Nup54 | 5ijo:F |
| | Nup58 | 5ijo:G |
| | Nup62 | 5ijo:H |
| Nup62-Nup58-Nup54 channel complex (repeat 2) | Nup54 | 5ijo:L |
| | Nup58 | 5ijo:M |
| | Nup62 | 5ijo:N |
| Nup62-Nup58-Nup54 channel complex (repeat 3) | Nup54 | 5ijo:R |
| | Nup58 | 5ijo:S |
| | Nup62 | 5ijo:T |
| Nup62-Nup58-Nup54 channel complex (repeat 4) | Nup54 | 5ijo:X |
| | Nup58 | 5ijo:Y |
| | Nup62 | 5ijo:Z |
| Connecting complex (repeat 1) | Nup155 | 5a9q:B |
| Connecting complex (repeat 2) | Nup155 | 5a9q:A |

(S10) Siegel, S. *Nonparametric statistics for the behavioral sciences.*; McGraw-Hill: New York, NY, US, 1956.

(S11) Pettersen, E. F.; Goddard, T. D.; Huang, C. C.; Meng, E. C.; Couch, G. S.; Croll, T. I.; Morris, J. H.; Ferrin, T. E. UCSF ChimeraX: Structure visualization for researchers, educators, and developers. *Protein Science* **2021**, *30*, 70–82.
